# Information regimes in population-graph neuroimaging: cohort context and implementation-dependent evaluation sensitivity

**DOI:** 10.64898/2026.10.02.756253

**Authors:** Zongqing Chen, Biantian Yu, Keliang Wan, Qian Zhang, Fanghong Zhang, Ning Zhong, Hongzhi Kuai

## Abstract

Population-graph models aggregate a query subject with an unlabeled cohort, so their reported performance can depend on the evaluation regime rather than on predictive information alone. Evaluation context has two separable axes: cohort visibility during *training* and the context available at *inference*. We study the training-time axis on an ABIDE-I cohort of 871 subjects (20 acquisition sites), comparing two population-graph implementations: a learned/adaptive graph (EV-GCN) and a static weighted graph (Parisot-GCN). Within each implementation the cohort, connectivity features, outer splits and training and selection rules are held fixed, and only the information available to the graph varies. The primary estimand is the implementation-by-training-context interaction, the difference between the P0-to-P1 AUC changes of the two implementations (ΔΔAUC). Across the two preselected outer splits (1024 and 2024), whose training and selection rules were fixed before the run, the direction is consistent and positive, with observed interaction range 0.06285–0.11967; the earlier audited split 666 shows the same direction and is reported as historical support, because its execution identity cannot be established to the same standard. Uncertainty is assessed within each split by paired subject and site-cluster bootstrap resampling; the intervals are conditional on the fixed partitions and the trained checkpoints. Splits are replication units: they are not pooled and no cross-split average is reported. These findings tie reported performance to the evaluation information regime in the two implementations compared, and motivate explicit reporting of cohort visibility.

## 1. Introduction

Population-graph models place subjects as nodes and propagate information between them, and were introduced for population-level disease prediction precisely so that non-imaging similarity between subjects could inform each subject’s prediction [1]. Successors replace the hand-crafted affinity graph with learned or adaptive graphs [2]. Because such models are trained on a graph that contains the query subjects themselves, the graph available at inference determines what the prediction for a query subject may depend on: whether other unlabeled test subjects are co-present, and whether those subjects come from the same acquisition site. A second, less often separated question is what the graph contained during *training*: whether the model was fitted on a graph that already contained the subjects it will later be queried on. The two axes are distinct, and conflating them makes reported performance estimates hard to compare.

Recent work has continued to expand this lineage. Population-level graph formulations have multiplied: a survey collects the current designs [3], along-side brain-to-population pipelines [4], heterophily-aware message passing [5], federated multi-site variants [6], cross-site individual–population integration [7] and knowledge-driven multi-graph constructions [8]. A second line broadens the setting itself: multi-site brain-network analysis framed as structure-aware domain adaptation [9] and self-supervised graph contrastive learning on resting-state fMRI [10]. These formulations differ in how the graph is built and how messages are aggregated, but the population-graph ones each keep the cohort inside the prediction path. Parallel graph-based work treated each subject’s connectivity network itself as the graph object and learned similarities between such graphs [11]; a recent benchmark evaluates each subject’s graph as an independent prediction unit on the combined ABIDE-I and ABIDE-II cohorts [12]. As this space widens, a reported performance figure depends not only on the learned representation but also on which cohort information the evaluation made available, and when.

Evaluation practice in this area is also known to be fragile. Leakage through feature selection and repeated subjects inflates connectome-model performance [13], and feature-selection leakage has been shown to invalidate reported autism fMRI accuracies [14]. Multi-site functional connectivity is confounded by site effects that limit generalisation [15], harmonisation of multi-site imaging data remains an active design question [16], and cross-validation design and sample size strongly affect the uncertainty of reported prediction estimates [17]. Population-graph evidence is usually reported under a transductive protocol. Inductive graph learning provides a general alternative to transductive node inference [18]; in population-GNN neuroimaging, a fully inductive single-subject inference protocol has recently been proposed as the deployment-relevant alternative [19]. A companion study from our group evaluates a site-aware population GNN in the same cohort, separates cohort-conditioned discrimination from performance at an unseen site, and traces a large part of its high transductive cohort accuracy to the same-site supervision available during learning [20]; that study manipulates the source of supervision and audits the evaluation as a whole, whereas the present study holds label supervision and inference context fixed and varies only which unlabeled subjects are visible to the graph during training. Our question is complementary to all of these: with the cohort, features, outer splits and training/selection rules held fixed within each implementation, how much do performance estimates change when only the *information available to the population graph* changes, and does that sensitivity depend on the implementation?

Read together, these strands motivate the contrast tested here: with label supervision and inference context held fixed, isolate the training-time visibility of unlabeled query nodes and measure whether its effect on reported performance differs between two controlled implementations. A population graph mixes two things that a conventional subject-level predictor keeps apart: a representation of the individual subject, and information about the cohort that subject is scored alongside. Which cohort information reaches the model is decided by the evaluation protocol, not by the data, so a reported performance figure is a property of the protocol as much as of the subject features; and the size of that protocol dependence may itself differ between implementations. The concern is not specific to autism or to either implementation: whenever subjects participate as nodes in a population graph, the evaluation protocol determines part of the information available to the predictor. We isolate this effect with information-regime interventions: the same implementations, data, splits and training rules, with the cohort information available to the graph as the only variable.

*Contributions.* (i) A two-axis taxonomy of population-graph evaluation regimes that separates training-time cohort visibility from inference-time query context and site familiarity (Table 1). (ii) Controlled cross-split evidence for an implementation-by-training-context interaction: two controlled implementations (EV-GCN, Parisot-GCN) evaluated on two primary preselected outer splits under one fixed cohort, one node-order convention, and training and selection rules locked before the run, with a third, earlier audited split reported as directional support. (iii) Separate reporting of split-level estimates, subject-bootstrap uncertainty and site-cluster uncertainty for the primary campaign; the earlier inference-context campaign’s estimands, resampling scheme and Holm-corrected framework are excluded from inference in this manuscript. (iv) A reusable reporting checklist for population-graph neuroimaging evaluation (Supplementary Checklist S1). The contribution is the controlled demonstration and the reporting discipline it implies, not a new predictor.

**Table 1:** Information available to the model in each evaluation regime. Rows P2–P5 are listed to define the taxonomy; they are not evaluated in this study and carry no result or inference here (evidence-scope column).

| Regime | Definition | Query nodes<br>in training<br>graph | Other<br>queries at<br>inference | Evaluation<br>role | Evidence<br>here |
| --- | --- | --- | --- | --- | --- |
| **P0 /**<br>**R0-T** | transductive | yes | yes | cohort-conditioned | evaluated |
| **P1 /**<br>**R1-C** | cohort-visible<br>inference | no | yes | cohort-conditioned | evaluated |
| P2 | batch /<br>context-dose<br>(k swept) | no | k of N | context-<br>budget<br>sensitivity | definition<br>only |
| P3 /<br>R2-Q | single-query<br>inference<br>setting | no | no | single-query<br>inference | definition<br>only |
| P4 | size-matched<br>target-site<br>context | no | site-matched | mechanism<br>probe | definition<br>only |
| P5 /<br>LOSO | trained on the<br>remaining<br>sites,<br>evaluated on<br>the held-out<br>site | no | held-out site | site generali-<br>sation | definition<br>only |

We make no claim of model superiority, no state-of-the-art ranking claim, and no new architecture.

## 2. Materials and methods

### 2.1. Cohort and connectivity

ABIDE-I derivatives are used under their original terms and are not redistributed: 871 subjects across 20 acquisition sites from ABIDE-I [21], following the canonical 871-subject cohort used by Abraham et al. [22]. The cohort comprises 403 autistic and 468 typically developing participants. Connectivity is taken from connectivity summarised on the CC200 parcellation [23]: one 200*×*200 Pearson correlation matrix per subject, not Fisher-*z* transformed in the controlled pathway, contributing the strict upper triangle as 19,900 features. All cohort quantities are read from the frozen cohort metadata record, which quotes the source record of each value. The primary campaign uses 5 outer mixed-site folds with disjoint fit/validation/query partitions, three split seeds and three training seeds (11, 22, 33), recorded in the two frozen split-registry tables listed in §2.5 and archived with the study.

### 2.2. Population-graph models and released-code characterisation

We distinguish two tracks. In **Track N** we inspect the released implementations to document how they construct data and select models; these observations are not used for any inference. **Track A** is the controlled evaluation reported here: disjoint fit/validation/query partitions, preprocessing fitted on the fit partition only, checkpoint and threshold selection on the validation partition only, and centralised context construction. Selection is fixed in the locked campaign definition: the checkpoint is the epoch with the highest validation AUC, evaluated every five epochs up to a 400-epoch budget (the final epoch is always evaluated), ties are broken towards the earliest epoch, and training stops after 20 validation checks without improvement. The decision threshold maximises Youden’s *J* on query-isolated validation predictions, with ties broken towards the value closest to 0.5 and then the lower one; nothing is refitted after selection. These rules were locked for the 1024 and 2024 extension runs; for split 666 they are not documented, and nothing in this section assigns them to that run retrospectively. Each Track-A implementation is checked against its released counterpart on fixed inputs. Released-code behaviour was characterised directly from the pinned upstream source and is summarised in the supplement; quantitative deviations observed during historical parity checks are not reported (see §6).

### 2.3. Information-regime taxonomy

Table 1 states, for each regime, which information is available to the model. The two axes are (a) training-time cohort visibility and (b) inference-time query context / site familiarity. Short names used throughout: P0/R0-T (test nodes present in the training graph), P1/R1-C (train-only graph, cohort-visible inference), P2 (context-dose, *k* swept), P3/R2-Q (single-query inference setting), P4 (size-matched target-site context) and P5/LOSO (site held out for training and evaluation, distinct from the site-omission analysis of fixed out-of-fold predictions used in this study). The *primary* campaign of this manuscript (S1) measures the training-context contrast P0 versus P1; the inference-context regimes P2, P3 and P4 belong to a separate, earlier query-context campaign, and P5 is a training-and-evaluation design rather than an analysis of existing predictions. All four are named here only to define the taxonomy; their outcomes are not part of the evidence reported in this manuscript. P2 was not part of S1. Figure 1 summarises the two axes of this taxonomy, the intervention that separates P0 from P1, and the scope of the replication design.

**Figure 1:**
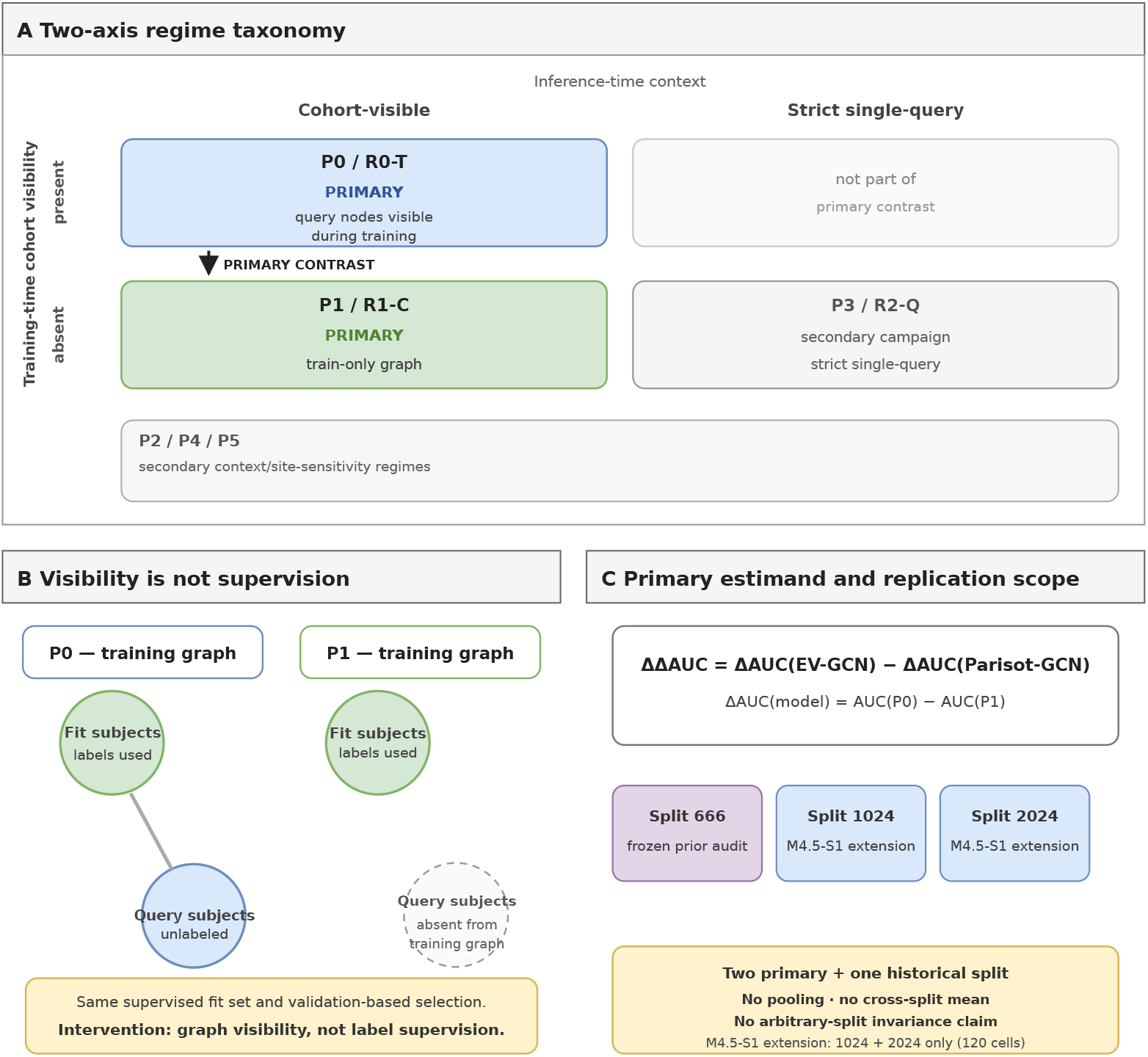
Information regimes and the primary controlled contrast. (A) Two-axis taxonomy separating training-time cohort visibility from inference-time query context. The primary P0-to-P1 intervention changes training-time graph visibility while retaining cohort-visible inference. (B) Query subjects are unlabeled in P0 and absent from the training graph in P1; supervised labels and validation-based selection are unchanged. (C) The primary implementation-by-training-context estimand and the two primary split-level replication units, with the earlier audited split shown separately as historical support; splits are not pooled. Secondary regimes vary inference-context size or site composition and are shown separately because they are not cells of the primary P0/P1 contrast.

### 2.4. Primary estimand

The primary estimand is the implementation-by-training-context interaction. Writing ΔAUC*_m_* = AUC*_m,P_*_0_ *−* AUC*_m,P_*_1_ for each implementation *m*, it is

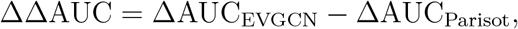

computed independently within each split from the three-seed probability ensemble of each condition. Positive values indicate a more negative P0-to-P1 AUC change for EV-GCN than for Parisot-GCN; they are not a claim that both implementations lose performance. The two implementations differ in architecture *and* in their model-specific graph operators, so the estimand is read at the level of the implementations compared here, not as a claim about architectures in general. Neither direction nor monotonicity is assumed, and the estimand is not a claim about which implementation is better in absolute terms.

*Visibility is not supervision.* In regime P0 the query subjects are present in the training graph as unlabeled nodes: their connectivity features and their graph relations are used, but their diagnostic labels are never used for optimisation. Regime P1 removes those query nodes from the training graph while keeping the supervised fit set, the validation-based checkpoint and threshold selection, and the training rules unchanged. The intervention therefore concerns cohort visibility in the graph, not label supervision, and this audit does not contrast a leaky pipeline with a clean one.

### 2.5. Preselected splits and replication design

The replication design uses three predefined mixed-site five-fold partitions of the same cohort, identified by their generation seeds 666, 1024 and 2024. Within each partition the five outer folds give disjoint fit, validation and query sets, and every subject is a query exactly once. Three quantities are kept apart throughout: the generation seed (666, 1024, 2024) labels a partition; the training seeds (11, 22, 33) are drawn within a partition; and the outer-fold index runs from 0 to 4 in the archived tables (folds are labelled 1–5 in the text and figures). The partitions are repeated splits of one cohort, not independent cohorts, and no result is reported as an independent replication of another. Membership of the partitions used here is recorded in two frozen tables archived with this study: `mixed_site_dx_5fold_seed666.csv` for seed 666, and `mixed_site_dx_5fold_v2.csv` for seeds 1024 and 2024, which also carries an unused reserve seed 2025; file hashes, field descriptions and the generation entry point are given in the supplement.

The reported partitions were not selected on the basis of their observed interaction. Split 666 comes from the earlier, independently audited campaign and precedes this analysis; the extension configuration, including the reserve seed, was frozen before any full controlled-graph result from that run existed. Splits 1024 and 2024 come from the later extension run, whose identifier is given with the reproducibility material; that run covers only those two splits: 120 cells in total, comprising 60 cells per split (two implementations *×* two regimes *×* 5 folds *×* three training seeds). Each split of the extension run provides exact-once query coverage of the cohort, as verified in the independent audit of that run; the earlier split-666 results carry their own audit and are not covered by the extension run’s identifier. Splits are treated as replication units; subjects recur across splits.

Evidence is stratified accordingly. Splits 1024 and 2024 are the primary controlled evidence: their training, selection and execution rules were fixed before the runs, and they supply the full 120-cell matrix. Split 666 is reported as historical directional support: its interaction is available, but its training and selection rules are not documented to the same standard and its condition-level AUCs are absent from the frozen source. The two roles are kept apart in the abstract, the results, Table 2, Figure 2 and the discussion; no estimate is pooled across roles, and the historical split is never counted as an independent cohort.

**Figure 2:**
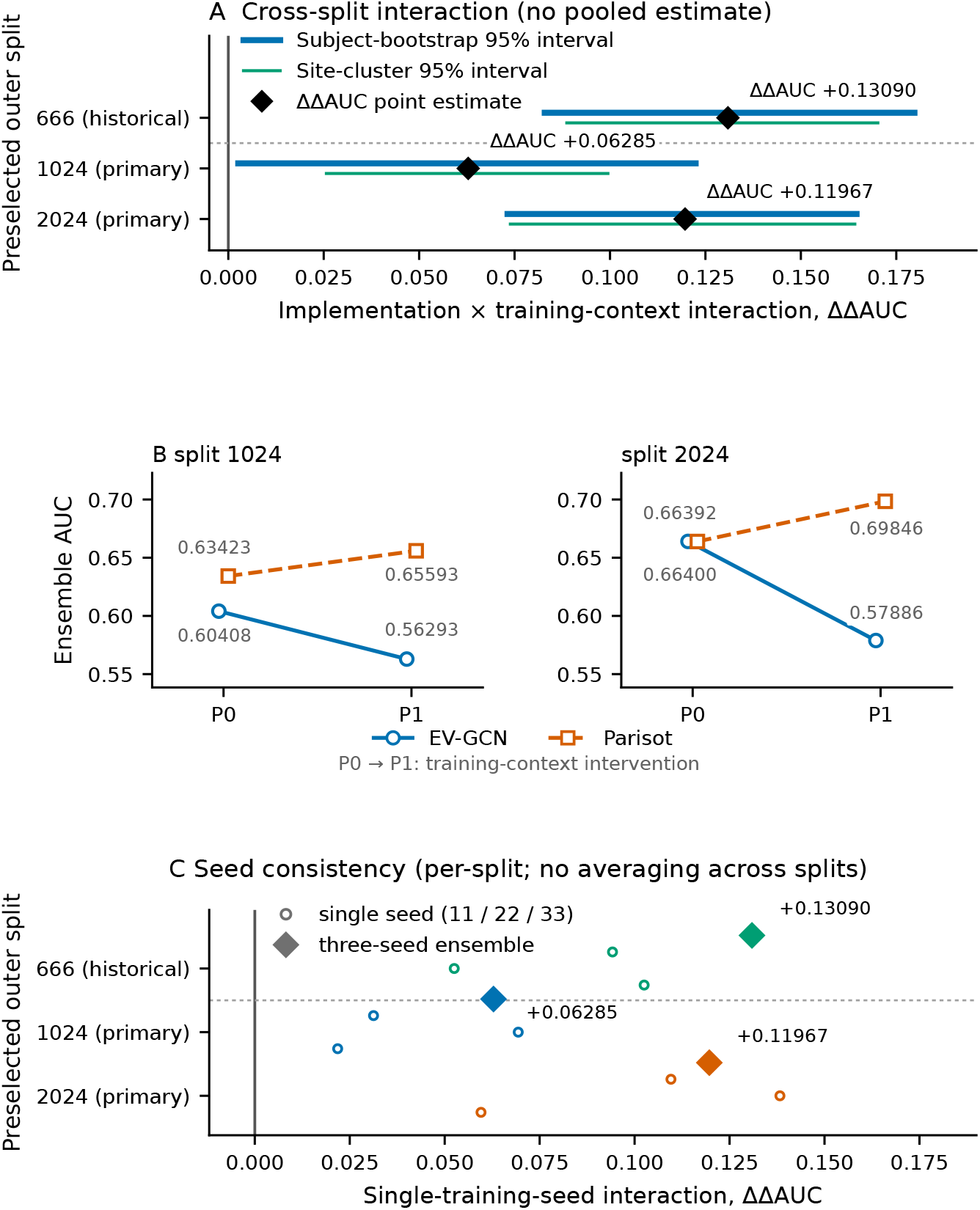
Primary interaction across the two primary splits (1024, 2024) and the earlier audited split (666), shown as historical support. (A) The implementation-by-training-context interaction for each split, with the subject-bootstrap interval and the site-cluster interval encoded separately; no pooled estimate is drawn. (B) The condition-level ensemble AUC before and after the intervention, for the splits whose condition-level AUCs are present in the frozen source; the prior audited split does not carry them and is omitted from panel B (it is shown in panels A and C). (C) The single-training-seed interaction values together with the three-seed ensemble, for each split. Within each regime, implementation markers are slightly offset horizontally for visibility; their vertical positions represent the reported AUC values.

**Table 2:**
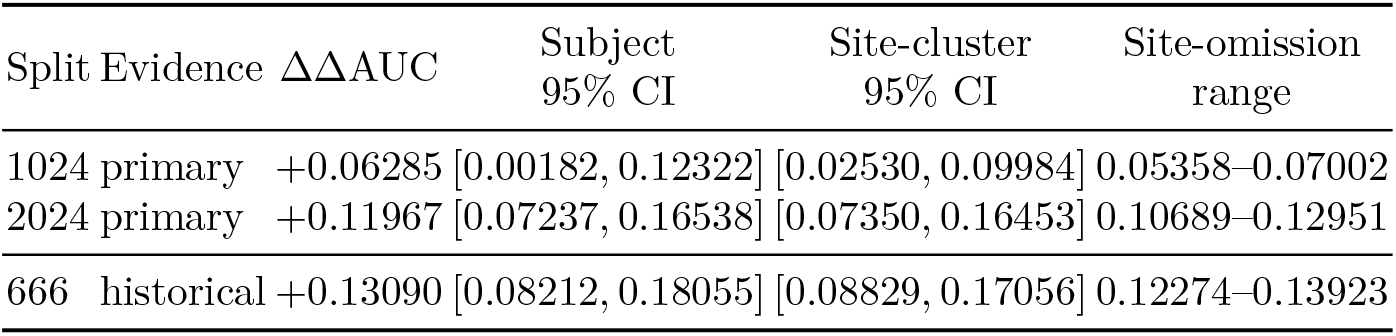
Cross-split replication of the implementation-by-training-context interaction over two primary splits (extension run, rules locked in advance) and one historical split (earlier audit, directional support only). Site-omission ranges omit one site at a time at evaluation, without refitting. Splits are replication units; they are not pooled.

### 2.6. Statistical inference

Primary inference is specified per split and never pooled across splits. For each split we report the three-seed ensemble interaction with (i) a paired subject bootstrap (10,000 replicates) that resamples subjects with the same resample used across the four conditions, and (ii) a paired site-cluster bootstrap (10,000 replicates) that resamples the acquisition sites and then subjects within drawn sites. Both interval types are two-sided 95% percentile intervals, taken at the 2.5th and 97.5th percentiles of the paired bootstrap distribution. Both are conditional on the frozen partitions and on the trained checkpoints, and both are marginal: they describe resampling of subjects and sites within a partition, are not adjusted for multiplicity across splits, and do not cover the variability of drawing new partitions or retraining. Site-omission (leave-one-site-out evaluation) estimates are reported as a sensitivity view of the same quantity, computed on the frozen out-of-fold predictions and not by refitting without the omitted site. Single-training-seed interactions are reported separately from the ensemble, because the ensemble averages per-subject probabilities within each condition before computing AUC and is therefore not the arithmetic mean of the seed-specific interactions.

#### Campaign separation

The earlier query-context campaign uses a different inferential framework (a per-subject context information score with a fixed smallest effect size of interest, and Holm-corrected contrast families) and a different protocol instance. The earlier campaign is historical background only: its estimands, resampling scheme and Holm-corrected framework are not merged into this manuscript’s protocol and do not apply to the split-specific interval estimates reported here.

### 2.7. Negative control for unintended context dependence

A feature-only control separates the regime effect from any unintended dependence of the data path on the training node set. The control reads the 19900 raw connectivity features, applies the same fit-fold feature selection (*k* = 2000) and row-sum normalisation as the controlled cells, and feeds the resulting 2000-dimensional vector to a two-layer perceptron (64 hidden units, no dropout). It has no graph, no message passing and no access to cohort statistics. Training uses Adam (learning rate 0.01, weight decay 5 *×* 10*^−^*^5^), full-batch updates with deterministic kernels, and at most 300 epochs; the validation AUC is evaluated at epoch 1, every five epochs thereafter and at the final epoch, the best checkpoint is kept with an improvement tolerance of 10*^−^*^9^, and training stops after 20 evaluations without improvement. The cached feature matrix has SHA-256 prefix 65925781adf5c8ae. The two regimes differ only in whether the query subjects are placed in the training batch as unlabeled rows, where they receive no loss: the training batch holds 609–787 subjects across folds. Because the control cannot use graph structure, the regimes must return the same predictions; the pre-registered tolerance is 1 *×* 10*^−^*^6^ on the absolute probability difference.

The control is an independent implementation of the feature path, not the released GNN code, and it differs from the controlled cells in three declared respects: the epoch budget is 300 rather than 400, the first epoch is evaluated as well, and no decision threshold is selected, because the control reports probabilities and AUC only. Its result therefore applies to this implementation and does not by itself certify the full population-graph training path.

### 2.8. Mechanism and secondary analyses

Further analyses were run in earlier stages of this project to probe the mechanisms behind regime sensitivity and to characterise the inference axis. Their outcomes are not part of the reported evidence (§6).

## 3. Results

### 3.1. Primary implementation-by-training-context interaction

The implementation-by-training-context interaction is positive on each of the two primary splits and, directionally, on the earlier audited split; the condition-level ensemble AUCs and the interaction for each split are reported in the supplement (Table S1). On splits 1024 and 2024 the interaction is +0.06285 and +0.11967 respectively; for split 666, which comes from the earlier audited run, the interaction is +0.13090. Condition-level AUCs for split 666 are not present in the frozen source, so that row reports the interaction only. On the two extension-run splits the underlying condition-level direction differs between the implementations: EV-GCN AUC decreases from P0 to P1 (0.60408 to 0.56293 on split 1024; 0.66400 to 0.57886 on split 2024), whereas Parisot-GCN AUC increases (0.63423 to 0.65593; 0.66392 to 0.69846). No common condition-level direction is therefore asserted across the two primary and the one historical split; the primary statement is the interaction, whose sign is the same in every split.

### 3.2. Replication across the two primary splits, with split 666 as historical support

The two extension-run splits carry the primary evidence: their point estimates are +0.06285 and +0.11967, with observed range 0.06285–0.11967, and the smaller split-1024 estimate has a subject-bootstrap interval whose lower bound is close to zero ([0.00182, 0.12322]). The earlier audited split contributes +0.13090, the same direction but a larger magnitude; because the training and selection rules of that campaign are not documented to the same standard and its condition-level AUCs are absent, we report it as historical directional support and do not average it with the primary splits. Both interval types and the site-omission range for each split are reported in Table 2. Site-omission estimates remain positive in every split, and the per-split ranges are drawn in the supplement (Figure S1). Subjects recur across splits, so the splits are not pooled and no cross-split average is reported; the supported statement is limited to consistency of direction and the observed range across the two primary splits, with split 666 as directional support. Figure 2 presents the same split-level results graphically, together with the condition-level response and the single-seed values.

### 3.3. Single-training-seed consistency

Every single-seed interaction is positive in every split, but their magnitudes vary, and the ensemble is not the arithmetic mean of the seed-specific values; the single-training-seed interactions alongside the three-seed ensemble are reported in the supplement (Table S2). Split-level replication therefore does not rest on one favourable training seed.

### 3.4. Negative control

Across the two primary splits, five folds and three training seeds (30 matched pairs, 60 independent runs), the feature-only control returned the same query predictions under P0 and P1: the largest absolute probability difference was 8.6 *×* 10*^−^*^11^, the mean 2.2 *×* 10*^−^*^12^, and the query-AUC difference was 0.0 in every pair. Re-running one configuration produced bitwise-identical output. The control carries real signal (median query AUC 0.72), so the agreement is not the degenerate case of a model that outputs a constant. Within this deterministic, feature-only implementation, changing the membership of the query rows in the training batch produced no probability difference above the registered tolerance; the result does not by itself certify the full population-graph training path.

### 3.5. Secondary analyses

Earlier mechanism and inference-context analyses are not part of the reported evidence (§6); the primary campaign does not depend on any claim about node-order behaviour.

## 4. Discussion

### Principal finding

Population-graph performance is not solely a property of subject-level features or model weights; it is also conditional on the information regime under which predictions are produced. Under a controlled protocol, removing training-time cohort context does not affect the two population-graph implementations equally: the implementation-by-training-context interaction is positive in both primary splits, with an observed interaction range of 0.06285–0.11967, and the earlier audited split is directionally consistent (+0.13090). The supported statement is that the controlled EV-GCN and Parisot implementations showed a consistently positive train-context interaction on the two primary splits, with split 666 as historical support; it does not establish invariance to arbitrary splits, nor isolate architecture from implementation-specific graph operators.

### Implementation-dependent regime sensitivity

The result is an interaction rather than a model-level superiority comparison. The effect of removing training-time cohort context differed between the two controlled implementations, yielding a positive interaction on each of the two primary splits and, directionally, on the historical split. On the two splits for which condition-level ensemble AUCs are available, EV-GCN AUC decreased from P0 to P1 while Parisot-GCN AUC increased; condition-level AUCs for the 666 split are not available in the stored results, so no common condition-level direction is asserted across the splits. The comparison is between two implementations that differ in architecture *and* in their model-specific graph operators, so we read the result as implementation-dependent regime sensitivity rather than as evidence about population-graph architectures in general. Reporting a single “transductive effect” for population graphs would hide this heterogeneity.

### Candidate mechanisms and how they could be tested

Three non-exclusive candidates are consistent with a larger interaction in the learned-edge implementation: (i) the learned edge function may itself depend on the training node set, so that removing query nodes changes the affinity structure; (ii) message passing over a sparser training graph changes degree normalisation and the effective neighbourhood size, which alters optimisation even when the affinity rule is fixed — a fixed-edge graph is therefore not immune to training-context changes by construction; and (iii) differences in capacity, depth and optimisation trajectory inherited from each implementation’s original design. These are candidates, not findings. A frozen-edge ablation, which holds the affinity at its P0 values and reruns the P1 training schedule with capacity, budget and normalisation fixed, would test (i) and (ii) as a change in the *visibility effect* rather than as a change in AUC alone. A subject-feature-only model, which cannot use cohort context, is the appropriate negative control for unintended context dependence in the controlled cells; its method is described in §2.7 and its result is reported in §3.4.

### Training context versus inference context

Our primary evidence concerns the training axis. Inference-time context (which other unlabeled subjects are present when a query is scored) is a separate axis measured by a different campaign with its own estimands; the two must be reported separately, because a model can be sensitive on one axis and not the other. P0 and P1 both retain cohort-visible inference: their contrast isolates training-time cohort visibility and does not establish performance under strict single-query deployment. The interaction is therefore not an estimate of prospective deployment overestimation.

### Implications for evaluation reporting

If a prediction is computed as *ŷ_i_* = *f* (*x_i_, X*_cohort_) where the cohort also participated in training, the reported performance answers a cohort-conditioned question. For prospective deployment this is a concrete reporting requirement: state the training-time cohort visibility, state the inference-time context, fix the node-order convention, and report the context composition. Neither regime is inherently invalid; rather, the regimes answer different evaluation and deployment questions. Recent work in medical-image analysis treats site and domain heterogeneity in multi-site brain networks as an explicit modeling problem [9]; the present results highlight a complementary issue: whether cohort context is available at all is itself part of the evaluation specification.

### Limitations

The evidence comes from a single dataset (ABIDE-I, fixed cohort of 871 subjects) with a heterogeneous site-size distribution. The primary result is established on two *preselected* splits, with a third, earlier audited split as directional support, and is not a claim of invariance to arbitrary splits; splits are not pooled and their intervals are not combined. Two controlled implementations are compared — selected for a matched information-regime intervention, not as an architecture benchmark — so the findings concern implementation-dependent sensitivity, and broader implementation coverage remains necessary before this interaction can be generalised across population-graph model families. Earlier mechanism, inference-context and historical-model analyses are not part of the reported evidence (§6); no claim here depends on them.

## 5. Conclusion

Population-graph prediction is not determined by an individual subject’s features alone: reported performance can also depend on which cohort information the evaluation protocol makes available. Under the controlled P0/P1 intervention, the two evaluated implementations differed in their sensitivity to training-time cohort context across both primary splits and, directionally, on the earlier audited split, so that sensitivity is implementation dependent rather than a property of population graphs in general. These findings motivate treating cohort visibility as an explicit component of population-graph evaluation and reporting rather than an implicit implementation detail, and tying claims about graph-based neuroimaging performance to the information regime under which the predictions are produced.

## 6. Reproducibility and data availability

ABIDE-I derivatives are not redistributed and are used under their original terms. Every primary number in this manuscript is generated from an archived, hash-identified results file whose entries carry, for each quantity, the exact text form and the source line of the stored evidence they were extracted from; the number layer and the result tables are generated from that file by a script, so no primary number is typed by hand. The released repository (https://github.com/ mindloop-lab/population-graph-information-regimes; release v1.0.1 archived at https://doi.org/10.5281/zenodo.23554740; v1.1.0, which adds the negative-control study and the revised checklist, at https://doi.org/10.5281/zenodo.23609463) contains the per-subject out-of-fold registry, the frozen split-membership tables under splits/, the figure generation scripts, the pinned upstream commits and the reporting checklist. The frozen split-membership tables are available under splits/ in the versioned software archive (v1.1.0; https://doi.org/10.5281/zenodo.23609463). Run identifiers, input provenance and the status of earlier numeric artefacts are recorded in the reproducibility supplement. The only earlier-run artefact that enters a reported result is the split-666 interaction already carried by the frozen record; the excluded historical parity deviations and mechanism analyses support no primary result or claim.

### Implementation and figures

Analysis and figure-generation code was drafted, revised and reviewed with AI assistance (ChatGPT/Codex, OpenAI, including GPT-6.1) under the authors’ direction, and Kimi K3 (Moonshot AI) assisted with organising the literature and with reviewing and polishing the Introduction; the authors verified the outputs against the frozen, hash-identified result sources. Figure 1 is a vector schematic rendered by a diagram script from its source drawing; Figure 2 and Figure S1 are generated by scripts from the archived result files; the graphical abstract is generated by a script from the same frozen values. No generative image model was used to create any figure.

### Items not reported

Quantities whose execution provenance could not be established to the standard required for the reported results are excluded from the evidence. This covers the historical parity deviations and the earlier mechanism and inference-context analyses; no primary result or claim depends on them, and the rule used to decide what may be reported is part of the released material (Supplementary Checklist S1).

## Declaration of generative AI and AI-assisted technologies in the manuscript preparation process

During the preparation of this work, the authors used ChatGPT/Codex (OpenAI, including GPT-6.1) to assist with language polishing, manuscript organisation and editorial revision, the drafting, revision and review of analysis and figure-generation code, and figure-layout refinement. Kimi K3 (Moonshot AI) was used to assist with organising the literature and reviewing and polishing the Introduction. These tools were used under the authors’ supervision. The authors reviewed and edited the AI-assisted material and verified the computational outputs against the frozen, hash-identified result sources. Reported numerical values were calculated or extracted by reproducible scripts from those sources; generative AI was not used to fabricate or alter research data or results. The authors retained responsibility for the study design, analytical decisions and final interpretation, and take full responsibility for the content of this publication.

## Declarations

### Ethics approval and consent to participate

This study was a secondary analysis of publicly available, de-identified data from the Autism Brain Imaging Data Exchange I (ABIDE-I). Ethics approval for the original data collection was obtained by each contributing site in accordance with its local institutional requirements; written informed consent was obtained by those sites from adult participants, or from the parents or legal guardians of minor participants, with assent obtained where applicable under the local protocols. No additional ethics approval was required for the present secondary analysis, and no participants were newly recruited.

### Consent for publication

Not applicable. This study used publicly available, de-identified data and contains no identifiable individual participant information.

### CRediT author contributions

Zongqing Chen: Conceptualization; Methodology; Investigation; Writing – original draft; Writing – review and editing; Project administration. Biantian Yu: Data curation; Investigation. Keliang Wan: Data curation; Investigation. Qian Zhang: Methodology; Writing – review and editing. Fanghong Zhang: Supervision; Funding acquisition; Writing – review and editing. Ning Zhong: Supervision; Methodology; Writing – review and editing. Hongzhi Kuai: Writing – review and editing.

## Appendix A. Reporting checklist

The reusable reporting checklist for population-graph evaluation is included as Supplementary Checklist S1 (v1, unchanged) and Checklist S2, which revises items 8, 16, 19 and 20: item 8 now applies only to a two-context comparison, and items 16, 19 and 20 admit both estimation-based and test-based inference. S2 carries the change record and the per-item applicability pointers; the study’s status under the v1 wording is unchanged.

