## Supplementary material for "Information regimes in population-graph neuroimaging: cohort context and implementation-dependent evaluation sensitivity"

This supplementary material provides the condition-level results, the seed-level detail and the site-omission sensitivity analysis supporting the main manuscript. It introduces no additional primary result or claim beyond those reported in the main text.

### 1. Condition-level results and seed-level detail

Table S1 reports the condition-level ensemble AUCs behind the primary interaction, for the two splits whose condition-level AUCs are present in the frozen source; the split from the earlier audited run carries the interaction but not the condition-level values. Table S2 reports the nine single-training-seed interactions and the three-seed ensemble that Figure 2C of the main text summarises. Both tables are the same generated inputs the main text reads, not copies.

Table S1: Condition-level ensemble AUC and the implementation-by-training-context interaction. AUCs are three-seed probability ensembles; the interaction is $(\mathrm{AUC}_{\mathrm{EVGCN},P0} - \mathrm{AUC}_{\mathrm{EVGCN},P1}) - (\mathrm{AUC}_{\mathrm{Parisot},P0} - \mathrm{AUC}_{\mathrm{Parisot},P1})$. Rows for splits 1024 and 2024 are the primary campaign; the row for split 666 is the earlier audited split, which carries the interaction only because its condition-level AUCs were not present in the frozen source.

| | EV-GCN | | Parisot-GCN | | |
|---|---|---|---|---|---|
| Split | $P0$ | $P1$ | $P0$ | $P1$ | ΔΔAUC |
| 666 | | | – | | +0.13090 |
| 1024 | 0.60408 | 0.56293 | 0.63423 | 0.65593 | +0.06285 |
| 2024 | 0.66400 | 0.57886 | 0.66392 | 0.69846 | +0.11967 |

Table S2: Seed-level consistency. Each cell is the single-training-seed interaction for that split; the ensemble column averages per-subject probabilities within each condition before computing AUC.

| Split | Seed 11 | Seed 22 | Seed 33 | Three-seed ensemble |
|---|---|---|---|---|
| 666 | +0.10251 | +0.05251 | +0.09418 | +0.13090 |
| 1024 | +0.02182 | +0.06936 | +0.03126 | +0.06285 |
| 2024 | +0.05956 | +0.13826 | +0.10957 | +0.11967 |

### 2. Released-code characterisation notes

In the released implementations, the static-graph model passes outer-test indices as validation indices and fits supervised feature selection on training indices; the learned/adaptive graph standardises edge inputs using complete-cohort statistics and evaluates test accuracy every epoch to select the saved checkpoint. These observations inform the controlled design but contribute to no inference in this paper.

### 3. Site-omission sensitivity

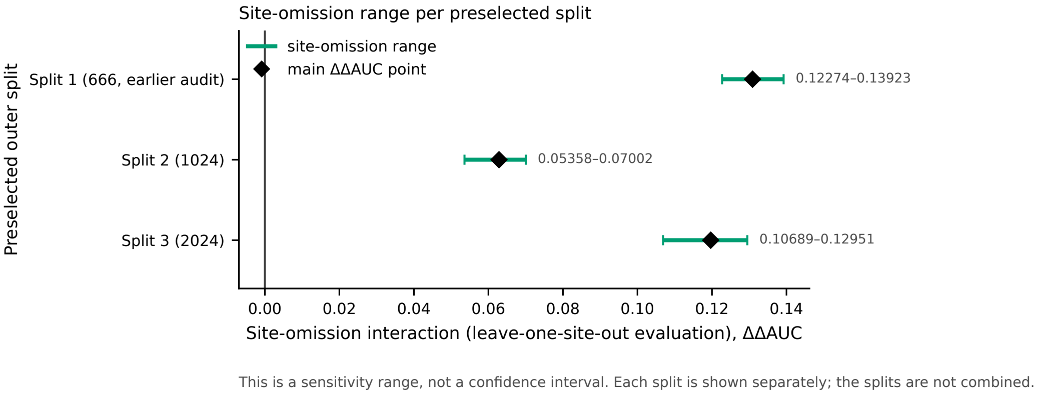


Figure S1: Leave-one-site-out sensitivity of the interaction, per preselected outer split. Each interval is the range of the interaction when one site at a time is omitted at evaluation from the frozen out-of-fold predictions (site-omission sensitivity, distinct from a leave-one-site-out training regime): a sensitivity range, not a confidence interval. Splits are replication units and are not pooled.

The main text reports the site-omission estimates as a sensitivity analysis (Section 2.6) and quotes the per-split range; Figure S1 shows the same ranges directly. Because subjects recur across splits, the three rows are not combined into a single summary.

### 4. Negative control

The control is a feature-only two-layer perceptron fed with the 19900 raw connectivity features after the same fit-fold selection ($k = 2000$) and row-sum normalisation used by the controlled cells; the network is 2000–64–2 with no dropout. It never reads a graph, an edge or a cohort statistic. Training uses Adam (learning rate 0.01, weight decay $5 \times 10^{-5}$), full-batch updates with deterministic kernels and at most 300 epochs, with the validation AUC evaluated at epoch 1, every five epochs and at the final epoch, a best- checkpoint improvement tolerance of $10^{-9}$, and early stopping after 20 evaluations without improvement. P0 and P1 differ only in whether the query subjects enter the training batch as unlabeled rows (no loss is assigned to them); the batch therefore holds 609–787 subjects. The pre-registered tolerance is $1 \times 10^{-6}$ on the absolute probability difference.

The control is an independent implementation of the feature path rather than the released GNN code, and it deviates from the controlled cells in three declared respects: a 300-epoch budget instead of 400, evaluation of the first epoch, and no decision threshold (the control reports probabilities and AUC only). The invariance result therefore applies to this implementation; it does not by itself certify the full population-graph training path.

Across 30 matched pairs (two splits, five folds, three training seeds; 60 separate processes, each written to its own file) the largest absolute probability difference was $8.6 \times 10^{-11}$ and the mean was $2.2 \times 10^{-12}$; the query-AUC difference was 0.0 in every pair, and a repeated run of one configuration was bitwise identical. The control is not degenerate: its median query AUC is 0.72. Raw inputs were cached once (SHA-256 prefix `65925781adf5c8ae`); the feature-selection and normalisation stage was recomputed inside each run, and the resulting matrix is fold-specific.

*Implementation caveat.* A stochastic variant (dropout 0.2) separates the two regimes, with a largest absolute probability difference of about $1.6 \times 10^{-1}$ and a query-AUC difference up to about $4.9 \times 10^{-3}$ (roughly five orders of magnitude above the tolerance). This is consistent with the number of dropout draws following the number of training rows rather than with any use of cohort information, and the same configuration re-run under one regime is bitwise identical; the cause was not isolated experimentally. The invariance verdict therefore rests on the deterministic variant, and a future invariance check on a stochastic model would need a random stream fixed per subject rather than per batch.

### 5. Split registry

The three partitions used in the main text are recorded in two frozen tables archived with this study: `mixed_site_dx_5fold_seed666.csv` and `mixed_site_dx_5fold_v2.csv`. Both are provided as supplementary material with this submission (under `splits_registry/`) and are archived in the software repository under `splits/` in release v1.0.1 (`https://doi.org/10.5281/zenodo.23554740`). They are reproduced unmodified from the campaign that produced them; nothing in this section regenerates them.

Table S3: Split registry. Rows are frozen membership tables; the two files are named in full in the text below. Hashes are SHA-256 over the archived file bytes.

| File | Generation seeds | Folds | Data rows | SHA-256 |
|---|---|---|---|---|
| seed-666 table | 666 | 5 | 4,355 | `fa689961...64b3` |
| extension table (v2) | 1024, 2024 (and the unused reserve 2025) | 5 | 13,065 | `d7d2a28c...af22d` |

Each row of a membership table has six fields: `subject_id`, `site_id`, `label`, `split_seed`, `outer_fold` and `role`. Within one (`split_seed`, `outer_fold`) cell the three roles `fit`, `validation` and `query` are disjoint, and every subject appears as `query` in exactly one of the five folds. `outer_fold` runs from 0 to 4 (folds are labelled 1–5 in the text). `label` is 1 for ASD and 0 for TD. The `split_seed` labels a partition; the training seeds 11, 22 and 33 are drawn inside a partition and are not recorded in these tables.

Partitions were built by stratification on site × diagnosis with five outer folds and a validation fraction of 0.125 of the working set. For the extension seeds this is confirmed by the documented generation entry point `scripts/generate_mixed_site_splits.py` driven by `configs/study/locked_campaign_v1.yaml`, which lists the split seeds $[1024, 2024, 2025]$ and the training seeds $[11, 22, 33]$; the reserve seed 2025 was not run. `scripts/generate_splits.py` is the earlier scikit-learn variant and is kept for historical reference. The seed-666 table predates this configuration: its run record states the same stratification design, but its execution identity (code and configuration) has not been recovered, so the table is reported by its own hash and is not attributed to either entry point.

### 6. Reporting checklist

Checklist S1: a reusable reporting checklist for population-graph neuroimaging evaluation. The 27 items below are reproduced from the archived checklist (v1, 2026-09-14) with their numbering, [must]/[if] conditions and wording unchanged; the artifact-pointer convention and the machine-readable copy are part of the released repository.

*A. Graph construction visibility.*

1. **[must]** Are the outer test subjects present as nodes in the graph used during training (parameter learning)? (`R0-T` vs `R1-C` distinction)
2. **[must]** Are outer test labels excluded from every fitting step (feature selection, normalization statistics, early stopping, threshold selection, hyperparameter search)?
3. **[if]** If test subjects are present at training time without labels, is this stated explicitly as *unlabeled cohort visibility* rather than *label leakage*?
4. **[must]** Are graph edges (or edge-candidate sets) constructed from the same subject set in training and inference? If not, name both sets.
5. **[must]** Are phenotypic variables of test subjects (e.g. acquisition site, sex, age) used in edge construction? Report yes/no per variable.

*B. Query context at inference.*

6. **[must]** Is a query subject's prediction computed with other unlabeled test subjects co-present in the inference graph (cohort-visible) or in isolation (single-query)?
7. **[if]** If cohort-visible, report the context size convention and whether it is the full remaining test cohort, a fixed batch size `k`, or sampled.
8. **[must]** Is the checkpoint identical between the cohort-visible and query-isolated runs? Report the checkpoint hash.
9. **[must]** Is the inference graph permutation invariant with respect to the query's node index? If not, state the fixed node-order convention and report the effect of the alternative convention.
10. **[must]** Is the decision threshold selected without using query labels?

*C. Site familiarity.*

11. **[must]** Report the acquisition-site composition of the inference context (same-site vs cross-site vs mixed) and, when possible, the size-matched same-site vs cross-site contrast.
12. **[if]** If leave-one-site-out (or unseen-site) results are reported, state that neither features nor labels of the held-out site were used at any fitting step.
13. **[must]** Do not describe a model that saw held-out-site features at training time as "unseen-site".

*D. Normalization and preprocessing scope.*

14. **[must]** Report the scope (fit set) of every normalization or harmonization step: full cohort before splitting, fold-local, or F-only.
15. **[if]** If any normalization statistic depends on the current inference graph (cohort-dependent normalization), report it as an information channel and, where possible, provide a frozen-statistics sensitivity run.

*E. Estimation and inference.*

16. **[must]** Report the primary estimand explicitly. If regime sensitivity is claimed, report a per-subject shift measure (e.g. mean $|\Delta \mathrm{p}|$) with its pre-specified smallest effect size of interest (SESOI).
17. **[must]** Report paired subject-level inference (bootstrap or permutation) using the same resample for both regimes.
18. **[must]** Cluster resampling by acquisition site when subjects are nested within sites (two-stage site$\rightarrow$subject), and compare with subject-level bootstrap as a sensitivity analysis.
19. **[must]** Pre-specify the families for multiple-comparison correction, and correct within family (e.g. Holm). Report raw and corrected p-values.
20. **[if]** If a model-by-regime interaction is claimed, report the difference-in-differences with a paired resample and a corrected p-value; a directional point estimate alone is not an interaction claim.
21. **[must]** Separate split-seed variance from subject-level sampling variance; never treat seeds or context draws as independent subjects.

*F. Reproducibility.*

**22. [must]** Provide the cohort manifest, split registry (with hash), model commit, environment lock, and per-cell checkpoint hashes.

**23. [must]** Provide per-subject out-of-fold predictions in a documented schema (subject, site, label, split seed, fold, train seed, model, regime, prediction), so that all reported numbers can be regenerated mechanically.

**24. [must]** State the node-order convention, the context sampling rule, and the threshold rule in the methods section, not only in code.

*G. Claims.*

**25. [must]** State the deployment assumption that the reported regime corresponds to. Cohort-visible inference answers a *cohort-conditioned* question; single-query inference answers a *prospective* question. Do not present either as the only correct protocol.

**26. [must]** Do not state that a model's absolute accuracy is *explained by* site composition unless a site-specific intervention supports that claim.

**27. [must]** Report the number of excluded sites/subjects and the exclusion rule.

*Applicability to this study*

Checklist S1 is reproduced verbatim (v1, 2026-09-14). The notes below state, for the items whose applicability depends on the study design, whether this study meets them; where it does not, the deviation is recorded rather than argued away.

*Checklist S2 — revision of items 8, 16, 19 and 20 (v2, 2026-10-22)*

Checklist S2 is the same 27-item instrument with four items revised: item 8 is made conditional, because it applies only when two inference contexts are compared, and items 16, 19 and 20 are rewritten so that an estimation-based analysis is not judged against a hypothesis-testing requirement. Checklist S1 is not withdrawn: the original items remain there, and this study's status under v1 (items 16, 19 and 20 not met) is unchanged and still recorded. The change record and a per-item applicability table with evidence pointers follow the items.

| Item | Requirement (abridged) | This study |
|---|---|---|
| 16 | Per-subject shift with a pre-specified SESOI, where regime sensitivity is claimed | **Not met.** The clause is triggered by a claim of regime sensitivity, which this study makes; the per-subject shift and SESOI it requires were not performed. What is reported is a split-level interaction with paired interval estimates, not a shift magnitude for individual subjects. |
| 19 | Pre-specify multiplicity families; report raw and corrected $p$ | **Not met.** No multiplicity family was pre-specified and no corrected $p$ is reported. Inference is interval-based; the per-split intervals are marginal and unadjusted, so they carry no simultaneous-coverage guarantee. |
| 20 | Interaction requires a paired resample and a corrected $p$ | **Partly met.** The interaction is estimated with a paired subject- and site-cluster bootstrap, separately per split; no $p$-value is reported and none is corrected, and the intervals are marginal rather than simultaneous. |
| 22 | Execution identity (code, configuration, selection rules) | Satisfied for 1024 and 2024 (locked campaign definition, checkpoint hashes). **Partial for 666**: its configuration and code identity were not recorded. |
| 23 | Per-subject out-of-fold predictions in a documented schema | Provided publicly for 1024 and 2024. **Not available for 666**, which predates the extension runs. |
| 24 | Explicit model-selection rules | Checkpoint metric, tie-break, early stopping and threshold rule are stated in §2.2 and fixed in the released configuration (`configs/study/`) for the 1024 and 2024 extension runs. They are **not documented for split 666** and are not attributed to it retroactively. |

*A. Graph construction visibility.*

1. **[must]** Are the outer test subjects present as nodes in the graph used during training (parameter learning)? (`R0-T` vs `R1-C` distinction)
2. **[must]** Are outer test labels excluded from every fitting step (feature selection, normalization statistics, early stopping, threshold selection, hyperparameter search)?
3. **[if]** If test subjects are present at training time without labels, is this stated explicitly as *unlabeled cohort visibility* rather than *label leakage*?
4. **[must]** Are graph edges (or edge-candidate sets) constructed from the same subject set in training and inference? If not, name both sets.
5. **[must]** Are phenotypic variables of test subjects (e.g. acquisition site, sex, age) used in edge construction? Report yes/no per variable.

*B. Query context at inference.*

6. **[must]** Is a query subject's prediction computed with other unlabeled test subjects co-present in the inference graph (cohort-visible) or in isolation (single-query)?
7. **[if]** If cohort-visible, report the context size convention and whether it is the full remaining test cohort, a fixed batch size `k`, or sampled.
8. **[if]** If two inference contexts are compared (for example the full remaining test cohort versus a single query), is the checkpoint identical between them, and is its hash reported?
9. **[must]** Is the inference graph permutation invariant with respect to the query's node index? If not, state the fixed node-order convention and report the effect of the alternative convention.
10. **[must]** Is the decision threshold selected without using query labels?

*C. Site familiarity.*

11. **[must]** Report the acquisition-site composition of the inference context (same-site vs cross-site vs mixed) and, when possible, the size-matched same-site vs cross-site contrast.
12. **[if]** If leave-one-site-out (or unseen-site) results are reported, state that neither features nor labels of the held-out site were used at any fitting step.
13. **[must]** Do not describe a model that saw held-out-site features at training time as "unseen-site".

*D. Normalization and preprocessing scope.*

**14. [must]** Report the scope (fit set) of every normalization or harmonization step: full cohort before splitting, fold-local, or F-only.

**15. [if]** If any normalization statistic depends on the current inference graph (cohort-dependent normalization), report it as an information channel and, where possible, provide a frozen-statistics sensitivity run.

*E. Estimation and inference.*

**16. [must]** State the primary estimand, its unit of analysis (subject, site, partition) and what would make the effect scientifically meaningful. If individual-level prediction shifts are part of the claim, report an appropriate shift measure and its aggregation rule. If a smallest effect size of interest is invoked, state its basis (substantive, prior evidence, or decision context) and the date it was fixed; a threshold derived from the current estimate is reported only as an exploratory sensitivity analysis.

**17. [must]** Report paired subject-level inference (bootstrap or permutation) using the same resample for both regimes.

**18. [must]** Cluster resampling by acquisition site when subjects are nested within sites (two-stage site→subject), and compare with subject-level bootstrap as a sensitivity analysis.

**19. [must]** State the scope of simultaneous inference. If the claim requires a family-wise or simultaneous statement, pre-specify the family and the correction (e.g. Holm) and report raw and adjusted quantities. If no simultaneous claim is made, say so explicitly and report intervals as marginal, per-partition intervals.

**20. [if]** If a model-by-regime interaction is claimed, estimate the difference-in-differences directly and report uncertainty that preserves the pairing (same resample for both regimes, and site clustering where subjects are nested in sites). If a hypothesis test is used, name the test, the null it samples under, and the multiplicity handling. A point estimate without any uncertainty statement is not an interaction claim; a point estimate with a paired interval suited to the estimand and to its dependence structure is.

**21. [must]** Separate split-seed variance from subject-level sampling variance; never treat seeds or context draws as independent subjects.

*F. Reproducibility.*

22. **[must]** Provide the cohort manifest, split registry (with hash), model commit, environment lock, and per-cell checkpoint hashes.
23. **[must]** Provide per-subject out-of-fold predictions in a documented schema (subject, site, label, split seed, fold, train seed, model, regime, prediction), so that all reported numbers can be regenerated mechanically.
24. **[must]** State the node-order convention, the context sampling rule, and the threshold rule in the methods section, not only in code.

*G. Claims.*

25. **[must]** State the deployment assumption that the reported regime corresponds to. Cohort-visible inference answers a *cohort-conditioned* question; single-query inference answers a *prospective* question. Do not present either as the only correct protocol.
26. **[must]** Do not state that a model's absolute accuracy is *explained by* site composition unless a site-specific intervention supports that claim.
27. **[must]** Report the number of excluded sites/subjects and the exclusion rule.

*Change record relative to v1*

Items 8, 16, 19 and 20 are revised. Item 8 becomes conditional ([if]) and is recorded as not applicable here, because no query-isolated inference contrast is reported; items 16, 19 and 20 are rewritten, and items 1–7, 9–15, 17–18 and 21–27 are unchanged in wording and numbering. The reason for the revisions is narrow: in v1, items 16, 19 and 20 treated one statistical route (hypothesis testing with a pre-specified SESOI and corrected $p$-values) as a universal requirement, so an estimation-based study could not satisfy them whatever its evidence; item 8 assumed a two-context comparison that a single-context study cannot answer. v2 keeps both statistical routes admissible and asks that the route be declared, and makes item 8 apply only when such a comparison is made. Under the v1 wording this study is still recorded as not meeting items 16, 19 and 20; that record is kept in Checklist S1.

*Applicability and evidence pointers, item by item*

Status is stated against v2. "Met (v2)" marks the three items whose assessment changes because the requirement changed; no other item changes status.

**1.** Met — §2.3; R0-T/R1-C defined

**2.** Met — §2.2 (fit-fold selection); OOF registry

**3.** Met — §2.3 terminology

**4.** Met — §2.2; split registry (§2.5)

**5.** Met — §2.2 (phenotype only in edge weights)

**6.** Met — §2.3; both regimes cohort-visible

**7.** Partial — inference context is the remaining test cohort (§2.3); the size convention is not stated in the main text

**8.** Not applicable — no query-isolated inference contrast is reported; the inference axis is excluded (§2.3)

**9.** Partial — fixed node order with the query last (§2.2); no permutation comparison is reported

**10.** Met — §2.2 threshold rule (Youden's $J$ on the validation partition only)

**11.** Partial — site composition of each split is in the registry; the inference-context composition is not tabulated

**12.** Not applicable — no held-out-site training regime reported

**13.** Met — no unseen-site claim

**14.** Met — §2.2

**15.** Not applicable — no cohort-dependent normalization

**16.** Met (v2) — §2.4 estimand; no per-subject shift claimed

**17.** Met — §2.6 paired bootstrap

**18.** Met — §2.6 site-cluster + subject bootstrap

**19.** Met (v2) — §2.6 marginal intervals declared

**20.** Met (v2) — §3.1–3.2 Delta-Delta AUC with paired intervals

**21.** Met — §2.6 seeds vs subjects

**22.** Partial — 1024/2024 yes; split 666 no

**23.** Partial — 1024/2024 yes; split 666 no

**24.** Partial — 1024/2024 yes; split 666 not documented

**25.** Met — §2.3 deployment assumption stated

**26.** Met — no site-composition explanation claimed

**27.** Partial — cohort size and site count are stated (§2.1); the exclusion rule is only in the released repository
